# Poincaré and SimBio: a versatile and extensible Python ecosystem for modeling systems

**DOI:** 10.1101/2024.01.10.574883

**Authors:** Mauro Silberberg, Henning Hermjakob, Rahuman S. Malik-Sheriff, Hernán E. Grecco

## Abstract

Chemical Reaction Networks (CRNs) play a pivotal role in diverse fields such as systems biology, biochemistry, chemical engineering, and epidemiology. High-level modelling of CRNs enables various simulation approaches, including deterministic and stochastic methods. However, existing Python tools for CRN modelling typically wrap external C/C++ libraries for modelling and simulation, limiting their extensibility and integration with the broader Python ecosystem. In response, we developed Poincaré and SimBio, two novel Python packages for the definition and simulation of dynamical systems and CRNs. Poincaré serves as a foundation for dynamical system modelling, while SimBio extends this functionality to CRNs, including support for the Systems Biology Markup Language (SBML). Poincaré and SimBio are developed as pure Python packages enabling users to easily extend their simulation capabilities by writing new or leveraging other Python packages. Moreover, this does not compromise the performance, as code can be Just-In-Time compiled with Numba. Our benchmark tests using curated models from the BioModels repository, demonstrate that these tools may provide a potentially superior performance advantage, compared to other existing tools. Additionally, to ensure a user-friendly experience, our packages use standard typed modern Python syntax that provides a seamless integration with Integrated Development Environments (IDEs). Python-centric approach significantly enhances code analysis, error detection, and refactoring capabilities, positioning Poincaré and SimBio as valuable tools for the modelling community.

## Introduction

Chemical Reaction Networks (CRNs) are a fundamental concept of modelling in numerous fields including systems biology, biochemistry, chemical engineering and epidemiology. They are comprised of a set of chemical species or biological entities and representation of complex interaction and transformation between them through a series reactions. These systems can be modelled and simulated through multiple approaches: deterministic Ordinary Differential Equations (ODEs) to model macroscopic behavior, Stochastic Differential Equations (SDEs) to model microscopic fluctuations, and jump processes (Gillespie-like simulations) to account for the discreteness of populations. Instead of directly writing the equations for each of these formulations which is error-prone and difficult to reuse, these models can be written in a higher-level description that can be compiled for these different types of simulations.

Several tools already exist for defining and simulating CRNs. BioSimulators.org (Shaikh et al. 2022), a registry of simulation tools, list at least 15 Python software including COPASI (Hoops et al. 2006), Tellurium (Choi et al. 2018) and PySB (Lopez et al. 2013). COPASI is a standalone software with a GUI for defining and simulating models. It’s widely used for its user-friendly interface and comprehensive features. There are also python bindings and interfaces for COPASI to allow advanced scripting. Tellurium is a Python-based modeling environment, however, it uses a C++ library called libRoadRunner in the backend to simulate models. PySB is a Python library that created a DSL using standard Python to define models which are then compiled to ODE using a Perl library called BioNetGen (L. A. Harris et al. 2016).

One limitation of these tools is that they are not extensible from Python, as they are not fully Python packages but wrap libraries in other languages to do the simulation. While they enable model definition and simulation control via Python scripts, they don’t fully leverage Python’s extensive package ecosystem. For example, in COPASI, users are restricted to predefined distributions for Random Parameter Scans and cannot utilize the diverse distributions available in `scipy.stats`. Similarly, Tellurium doesn’t allow the use of Python solvers, as adding new integrators requires C++ implementation.

Another challenge is the way models are written. Many tools use a Domain Specific Language (DSL) for coding, and support the System Biology Markup Language (SBML) as an exchange format (Hucka et al., n.d.), as direct SBML coding is impractical. A DSL allows reuse of code in different programming environments, but it will not be recognized by default in Integrated Development Environments (IDEs). Therefore DSLs cannot provide the development experience such as syntax highlighting, code completion, refactoring, and static analysis, unless such support is specifically developed. Tellurium uses a Domain Specific Language (DSL) called Antimony (Smith et al. 2009), which can be translated to SBML. An extension for Visual Studio Code was developed, but its maintenance could be demanding task for the systems biology community. PySB, using Python’s dynamic nature, created a DSL within Python. By default, it uses global state to create species and parameters, without assigning them to Python variables or adding them explicitly to the model, but this approach is not fully compatible with IDEs, affecting the development experience.

To overcome these limitations, we developed poincaré and SimBio, open-source Python packages for defining and simulating systems. Poincaré allows defining differential equation systems, while SimBio builds on it for defining reaction networks. They are focused on providing an ergonomic experience to end-users by integrating well with IDEs and static analysis tools through the use of standard modern Python syntax. Since they are coded in Python, every part from model definition to simulation can be extended from Python syntax. Being the first-ever pure Python packages for systems modelling, they offer extensive extensibility, from simple tasks like reusing integrators defined in other packages, to complex ones like altering the compilation process to leverage some structure in the equations. For example, using a for-loop in the compiled equations could improve the runtime performance if there is some repetitive structure in the system, as happens in spatial modelling. The models built using these packages can be introspected to create other representations, such as graphs connecting species and/or reactions, or tables with parameters or equations. Furthermore, they have a modular architecture with a clear separation of concerns, making it easier to maintain or to contribute new code, which is beneficial for developers and maintainers. We showcased the reliability of these tools by benchmarking them against the simulation results from other tools. We also highlighted the substantial performance improvements our tools offer, as this is crucial for construction and simulation of models of whole cells and organisms, which necessitate the simulation of significantly large-scale models.

## Results

Modular code architecture makes code reusable, extensible, and easier to maintain. Therefore, we split the code to define and simulate reaction systems into three Python packages: symbolite, to create symbolic expressions; poincaré, to define dynamical systems; and simbio, to define reaction systems and interface with systems biology standards such as SBML. These are pure Python packages with standard dependencies from the PyData scientific stack such as NumPy (C. R. Harris et al. 2020) and pandas (McKinney 2010). They are published in PyPI (the Python Package Index), where links to the source code and documentation hosted in GitHub can be found, and can be easily installed with `pip install <package_name>`.

Symbolite is a lightweight symbolics package to create algebraic mathematical expressions. Symbolite expressions can be inspected and compiled to various backends. Currently, we have implementations for NumPy (C. R. Harris et al. 2020); Numba (Lam, Pitrou, and Seibert 2015), a Just-in-Time (JIT) compiler to LLVM; SymPy (Meurer et al. 2017), a library for symbolic mathematics; and JAX (Bradbury et al. 2018), a library that support automatic differentiation and compilation to GPUs and TPUs. Symbolite is designed to facilitate the easy integration of new backends.

### Versatile modelling and simulation of dynamical systems with Poincaré

Poincare is a package to define and simulate dynamical systems. It provides a `System` class, where one can define `Constants, Parameters, Variables`, and create equations linking a variable’s derivative with an expression (Figure 1a). It also allows to define higher-order systems by assigning an initial condition to a `Derivative` (Figure 1b).

**Figure 1:**
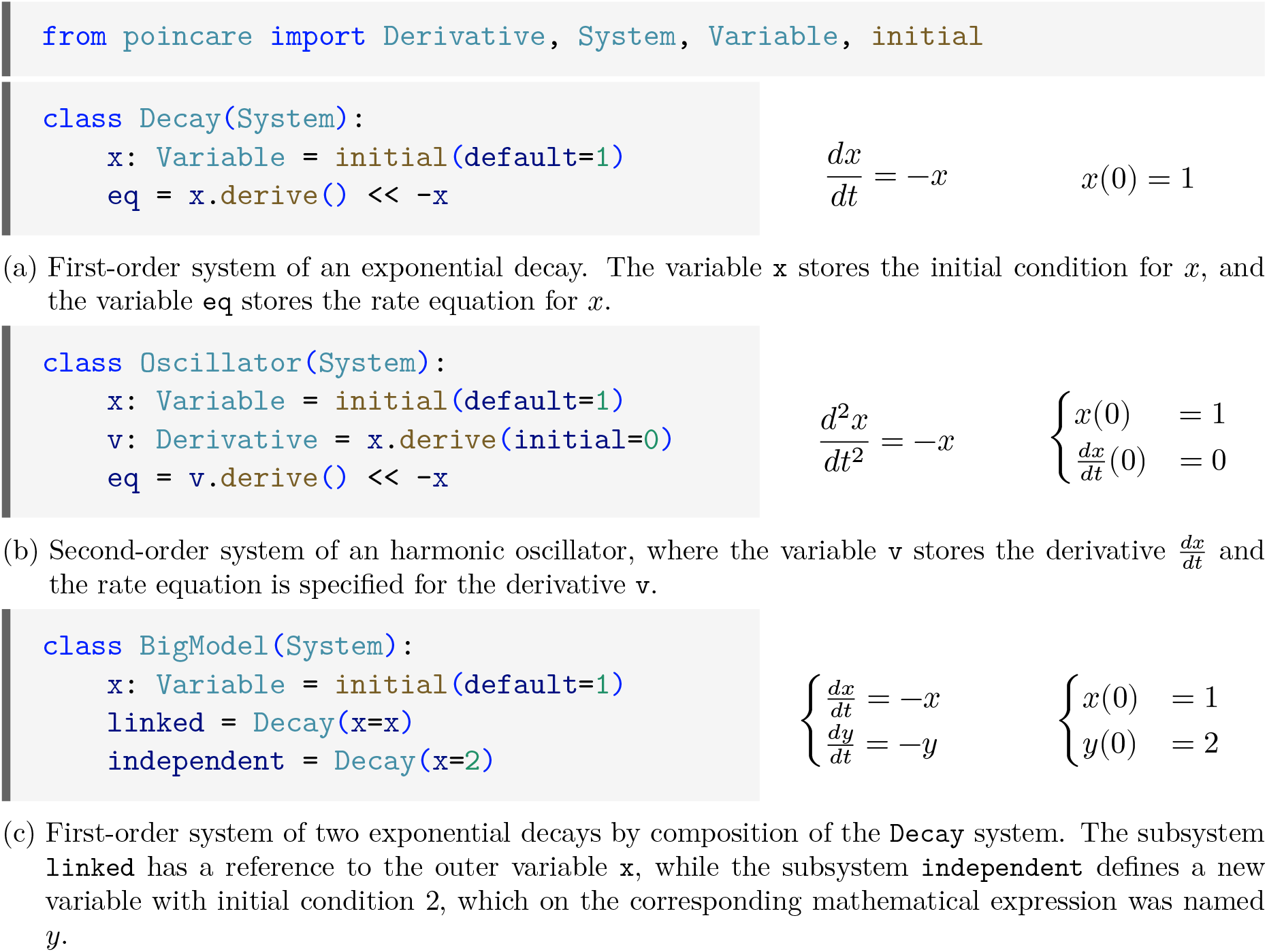
Code and corresponding mathematical expressions for different systems.

Utilizing classes for system definition offers several advantages:

1. The variable name to which a component is assigned can be automatically saved in the component for introspection (i.e., `Oscilator.x.name == “x”`),
2. It provides a namespace such that allows to easily define of multiple independent models in the same script,
3. It allows IDEs to provide autocomplete and refactoring capabilities (Oscillator.<TAB> shows x, v and eq),
4. It allows creation of instances which can be composed into a bigger model (Figure 1c).

For this last point, IDEs that support `dataclass_transform` (De Bonte and Traut 2021) can provide a tooltip with the expected signature (Figure 1c). This requires the use of type annotations which play a more significant role in static type checking, as they can help to identify errors before running the code. For instance, to parameterize the initial conditions of variables we have to use a `Constant`. If we try to use a `Parameter`, which could be a time-dependent expression, it is flagged as a type error (Figure 2).

**Figure 2:**
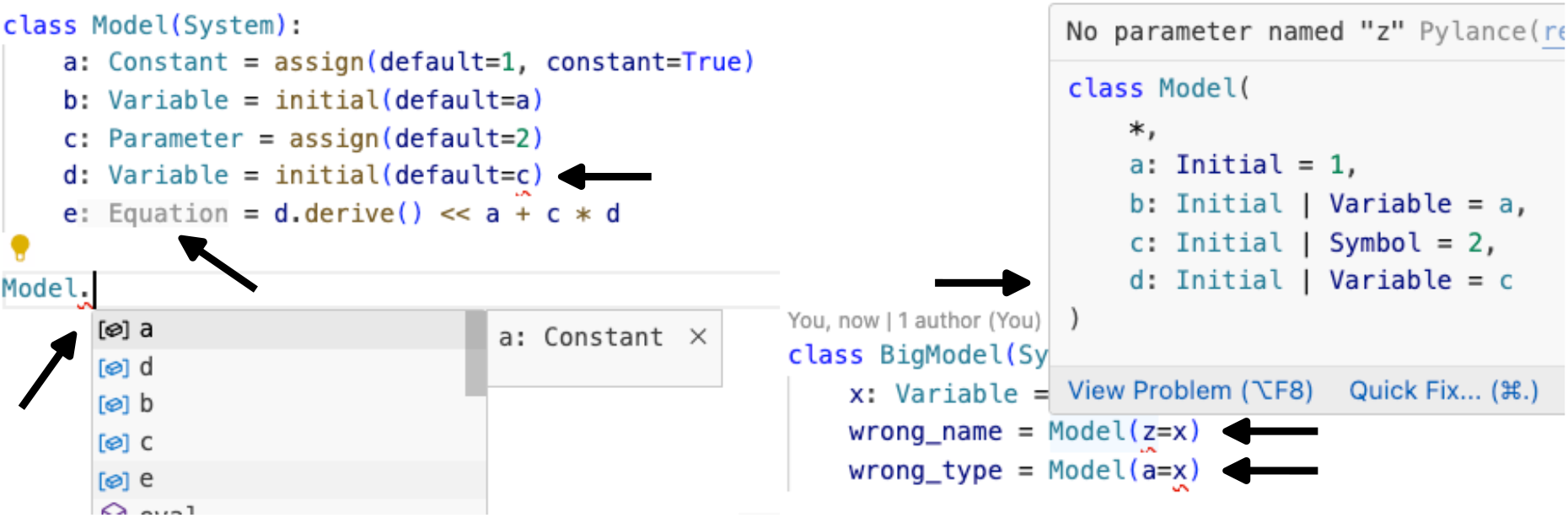
Screenshots of Visual Studio Code showing tooltips and highlighted type errors. `Constant` are assigned with `assign`(…, `constant=True`) and can be used to link `Variables` initial conditions, while trying to use a `Parameter` instead is flagged as type errors (red underlining). The IDE automatically recognizes `e` as an `Equation`, and provides autocompletion of the variables. A tooltip is shown when composing models, which show the expected variables and their default values. The IDE highlights wrong names (`z` is not a name in `Model`) and mismatched types (`x` is `Variable` and `a` must be a number or a `Constant`)

To simulate a system, we created a `Simulator` instance (Figure 3), which compiles a given system and interfaces with solvers. By default, it creates a first-order ordinary differential equation (ODE) system using `numpy` as a backend. This can be easily switched to other solvers. The Simulator wraps the output in a `pandas.DataFrame`, which can be easily plotted with the standard `plot` method.

**Figure 3:**
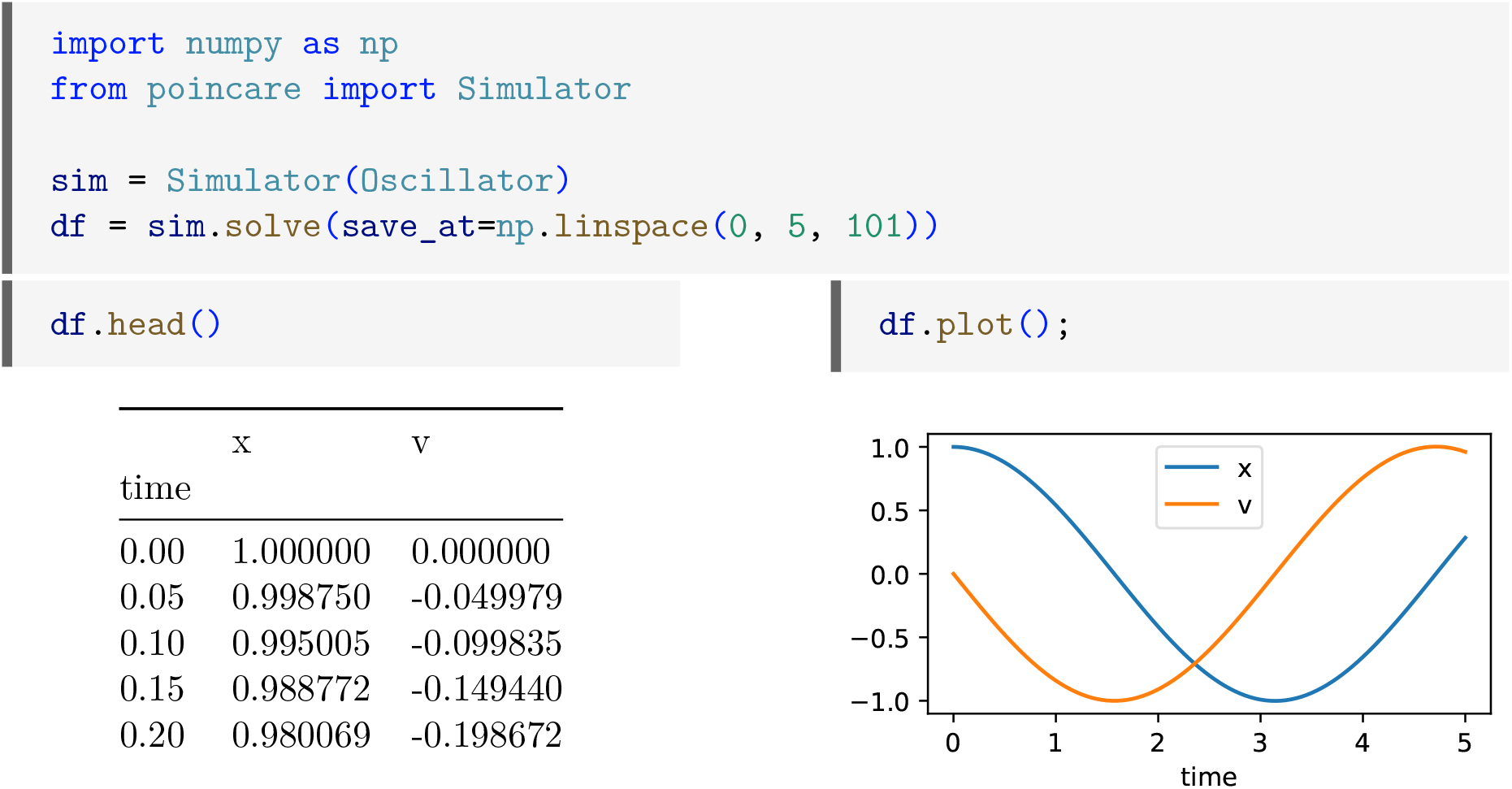
Simulation of the `Oscillator` system from Figure 1b. The output is a `pandas.DataFrame` with a column for each variable and the time as index. It is inspected and plotted with the `pandas` methods.

Switching backends to “numba” can improve model runtime on a factor up to x30 or more for big or long running models, however incurs in a compile time penalty for the first run which must be taken into account.

### Extensible definition of reaction networks using SimBio

For the reaction networks, our focus is on first-order differential equations that describe the rate of change of species. SimBio simplifies the definition of these network models by introducing `Species`, which consists of a `poincare.Variable` and a stoichiometric number, and RateLaw, a construct that converts reactants into products taking into account the stoichiometry (Figure 4). Additionally, SimBio features `MassAction`, a subclass of `RateLaw`, which intuitively incorporates reactants into the rate law (Figure 4).

**Figure 4:**
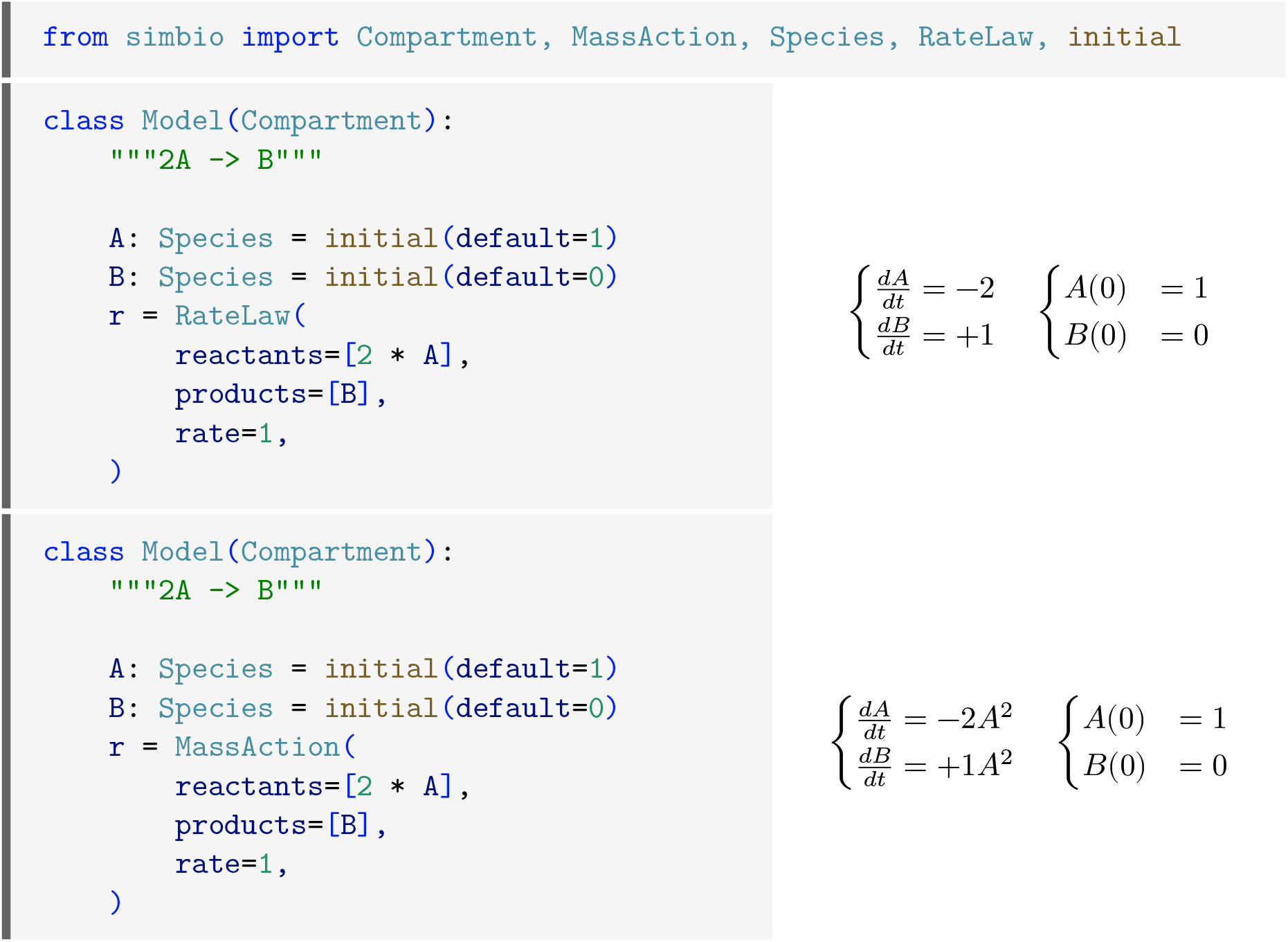
A reaction system for species *A* and *B* with initial conditions 1 and 0, respectively. A single reaction transforming 2*A* into *B* is saved in variable r. The rate 1 is specified directly for `RateLaw`, and is proportional to the reactants for `MassAction`.

Several commonly used reactions are predefined as MassAction subclasses, such as MichaelisMenten (*S* + *E* ↔ *ES* → *P* + *E*) and its approximate form without the intermediate species *ES*, and it is also simple to implement your own as subclasses of RateLaw or MassAction. Additionally, SimBio supports importing from and exporting to SBML, and downloading them directly from BioModels (Malik-Sheriff et al. 2020) (Figure 5).

**Figure 5:**
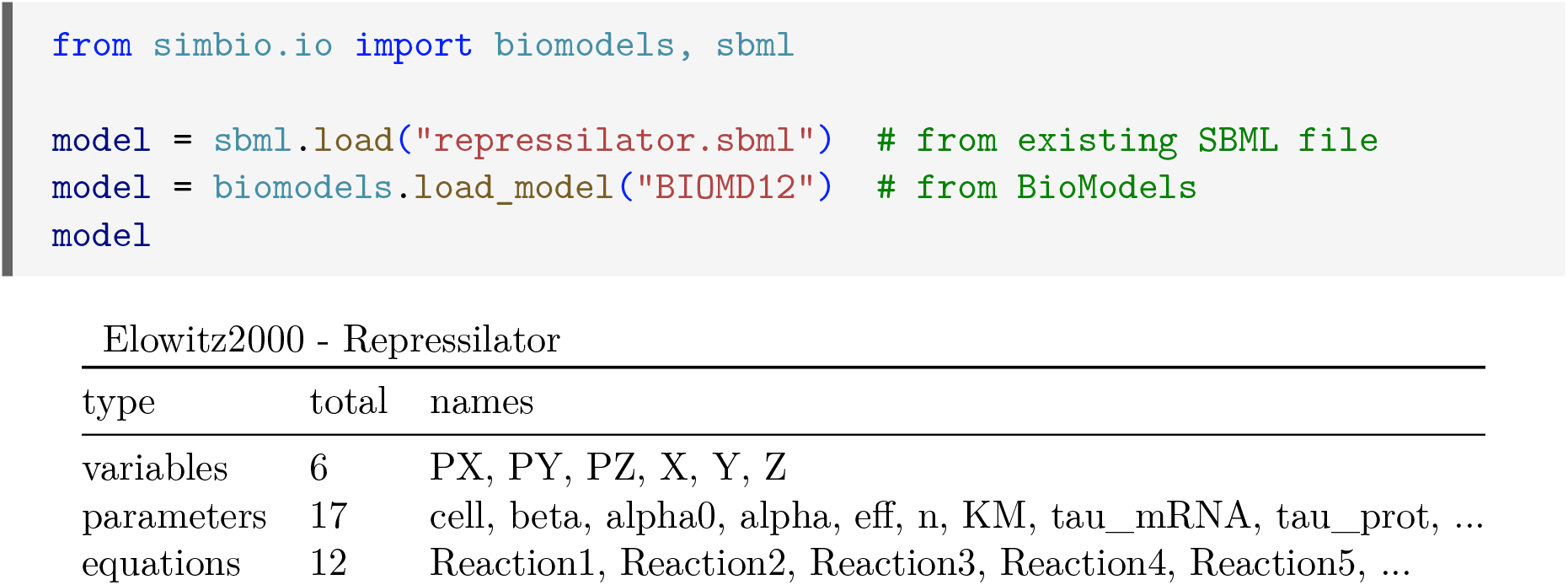
Creation of a model from a local SBML file or one uploaded to BioModels.

### Reproducibility and performance

To evaluate SimBio’s reproducibility, we analyzed curated SBML models from BioModels (Malik-Sheriff et al. 2020). Among the first 100 curated models from BioModels we selected 60 which did not contains events, as SimBio doesn’t support events yet. We simulated the selected models with COPASI and used the simulated results as ground truth, and demonstrated that SimBio reproduces the results with minor numerical differences depending on the solver tolerance.

For performance testing, we ran simulations using COPASI, Tellurium/RoadRunner, and SimBio. Within SimBio, both NumPy and Numba backends were considered. The LSODA solver was used for COPASI and SimBio, while for Tellurium the comparable CVODE solver was used. In all cases, we used relative and absolute tolerances of 10^−6^. We measured three simulation stages: loading, the initial (cold) run, and subsequent (warm) runs for each model.

For COPASI and Tellurium/RoadRunner, we noted that its runtime depended on the number of evaluation points, something that does not seem to happen with SimBio (Figure 6, left). While SimBio’s NumPy backend is slower than both COPASI and RoadRunner, we obtained an order of magnitude speed-up using the numba backend putting it on par with them. A user might have to consider the trade-off between compilation and run times, as the compilation of the right-hand-side (RHS) code might take longer than the runtime itself, and not be worth it for running the model only once. Another speed-up in the runtime can be had by switching the LSODA scipy solver for a more efficient `numbalsoda` implementation, which avoids the Python interpreter between each of the integration steps. This last combination beats all other methods, which is also true for the other models we tested (Figure 6, right).

**Figure 6:**
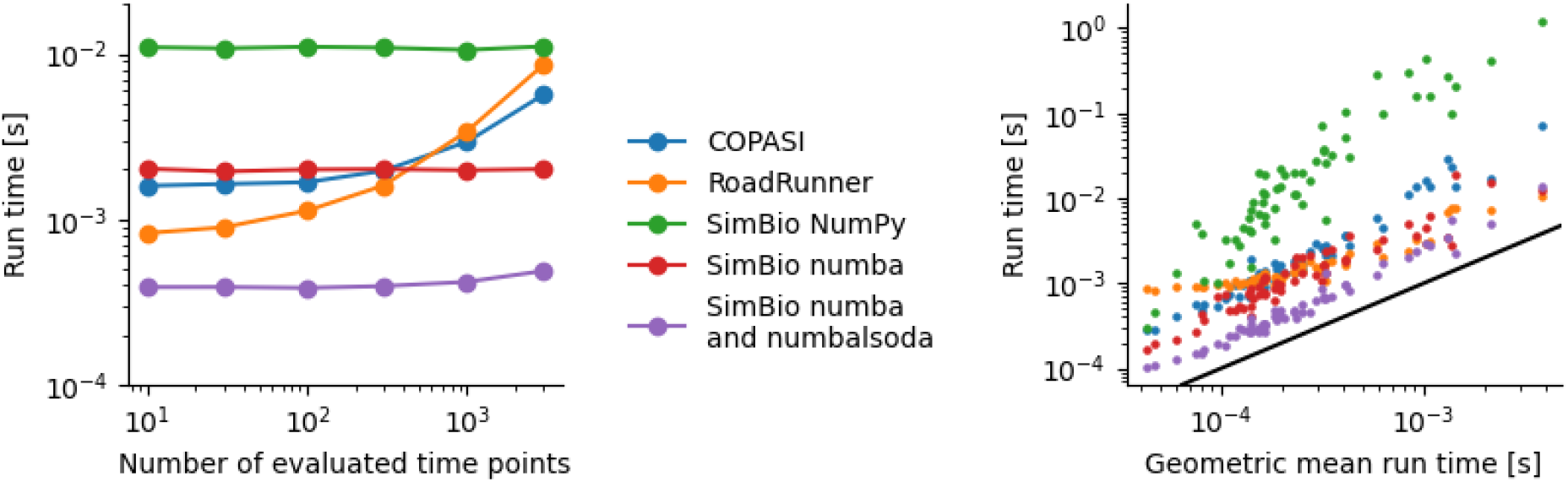
Performance of different softwares to solve models from the curated section of BioModels. (left) Run time for the model BIOMD3 as a function of the number of output points. (right) Run time for different models for 300 output points, using the geometric mean of the different softwares to order them.

## Discussion

In this article, we introduced a suite of Python packages we developed for defining and simulating dynamical systems and reaction networks. These packages are deeply integrated with Integrated Development Environments (IDEs), enabling code analysis tools to identify errors prior to execution and assist in refactoring and code completion. We adopted standard modern Python syntax to ensure seamless IDE integration, supported by the extensive Python community.

Our approach differs from previous tools in that both the model definition and its compilation into an Ordinary Differential Equation (ODE) function are entirely Python-based. This approach simplifies the development of various simulation methods, including performance enhancements that exploit specific model structures. Importantly, being Python-based does not compromise performance compared to C/C++ tools, as the ODE functions can be Just-In-Time (JIT) compiled using Numba.

The inclusion of SBML support facilitates the effortless reuse of models created by the systems biology community, along with the vast collection of public models hosted in the BioModels repository. The modular architecture of these packages facilitates their reuse, enhancement, and extension by the wider Python community. For instance, an individual from outside the systems biology field could contribute a stochastic integrator to poincaré, which would then be available in SimBio. This clear separation of concerns also makes the packages more comprehensible, lowering the barrier for contributing improvements or new features. Such an architecture ensures their maintainability and ongoing development well into the future.

## Resources

Source code repositories for these packages can be found at https://github.com/maurosilber/poincare and https://github.com/hgrecco/simbio.

